# Conserved Circuits for Direction Selectivity in the Primate Retina

**DOI:** 10.1101/2021.07.21.453225

**Authors:** Sara S. Patterson, Briyana N. Bembry, Marcus A. Mazzeferri, Maureen Neitz, Fred Rieke, Robijanto Soetedjo, Jay Neitz

**Affiliations:** Center for Visual Science, University of Rochester, Rochester, NY, 14620; Department of Ophthalmology, University of Washington, Seattle, WA, 98109; Department of Physiology and Biophysics, University of Washington, Seattle, WA, 98195; Washington National Primate Research Center, University of Washington, Seattle, WA, 98195

## Abstract

The detection of motion direction is a fundamental visual function and a classic model for neural computation^1,2^. In the non-primate mammalian retina, direction selectivity arises in starburst amacrine cell (SAC) dendrites, which provide selective inhibition to ON and ON-OFF direction selective retinal ganglion cells (dsRGCs)^3,4^. While SACs are present in primates^5^, their connectivity is unknown and the existence of primate dsRGCs remains an open question. Here we present a connectomic reconstruction of the primate ON SAC circuit from a serial electron microscopy volume of macaque central retina. We show that the structural basis for the SAC’s ability to compute and confer directional selectivity on post-synaptic RGCs^6^ is conserved in primates and that SACs selectively target a single ganglion cell type, a candidate homolog to the mammalian ON-sustained dsRGCs that project to the accessory optic system and contribute to gaze-stabilizing reflexes^7,8^. These results indicate that the capacity to compute motion direction is present in the retina, far earlier in the primate visual system than classically thought, and they shed light on the distinguishing features of primate motion processing by revealing the extent to which ancestral motion circuits are conserved.

---

Neurons responding preferentially to motion in specific directions are found across species and throughout the visual system^1,2^. The underlying mechanisms have been extensively studied in ON and ON-OFF dsRGCs of the non-primate mammalian retina^9^. Each consists of multiple subtypes preferring motion in different directions^10^. Their direction selectivity begins with SACs, radially-symmetric interneurons present in every mammalian retina studied to date^5,11^. SAC dendrites operate independently, computing outward motion from the soma^12^ and providing selective GABAergic inhibition to dsRGC subtypes preferring motion in the opposite direction^3,4^.

The intensive study of direction selective retinal circuits has yielded significant insight into the general principles of neuronal computation^1^, yet the direct applications to primate vision are unclear. Despite being a standard feature in the early visual systems of other species, direction selectivity has yet to be demonstrated in the primate retina. Several lines of evidence indicate some retinal capacity to compute motion direction may be conserved^13–15^, yet primate dsRGCs remain elusive. As such, a classic interpretation is that the expanded primate cortex replaced the need for retinal direction selectivity and the other highly-specialized computations found in non-primate retinal ganglion cells (RGCs)^16,17^. Alternatively, the absence of primate dsRGCs from the literature could reflect a sampling bias. The primate retina is dominated by three RGC types and the rarity of the other ~15 anatomically-defined types severely limits the possibility of identifying dsRGCs with the electrophysiology approaches used in other species^9,18^. Thus, the underlying question is not only whether primate dsRGCs exist, but also how to find them.

An alternative strategy is to identify candidate dsRGCs from the neurons post-synaptic to SACs. However, this approach is currently limited by a lack of information on primate SAC circuitry. Here we use serial block-face scanning electron microscopy and connectomics^19^ to fill this gap in knowledge and determine the extent to which the mammalian retinal direction selectivity circuitry is conserved in primates.

## Connectomic Reconstruction of ON SACs

We reconstructed a population of 8 ON SACs from a 220 x 220 x 170 μm volume of macaque central retina (~1.5 mm inferior to the fovea) sectioned horizontally at 90 nm and imaged at 7.5 nm/pixel (see **Methods**)^20^. The volume spanned from the photoreceptor outer segments to the ganglion cell layer, enabling 3D reconstruction of complete retinal circuits while maintaining the resolution necessary to identify synapses. Each reconstructed ON SAC exhibited the stereotyped morphological features and stratification described across species (**Fig. 1**)^5^. We focused on the ON SACs because primate OFF SACs are nearly absent from the fovea and outnumbered 10:1 by ON SACs in the periphery^5,21^. Consistent with previous reports of primate SACs, the dendritic fields in **Fig. 1** are sparse compared to non-primate SACs, both in branching density and coverage factor5.

**Figure 1.**
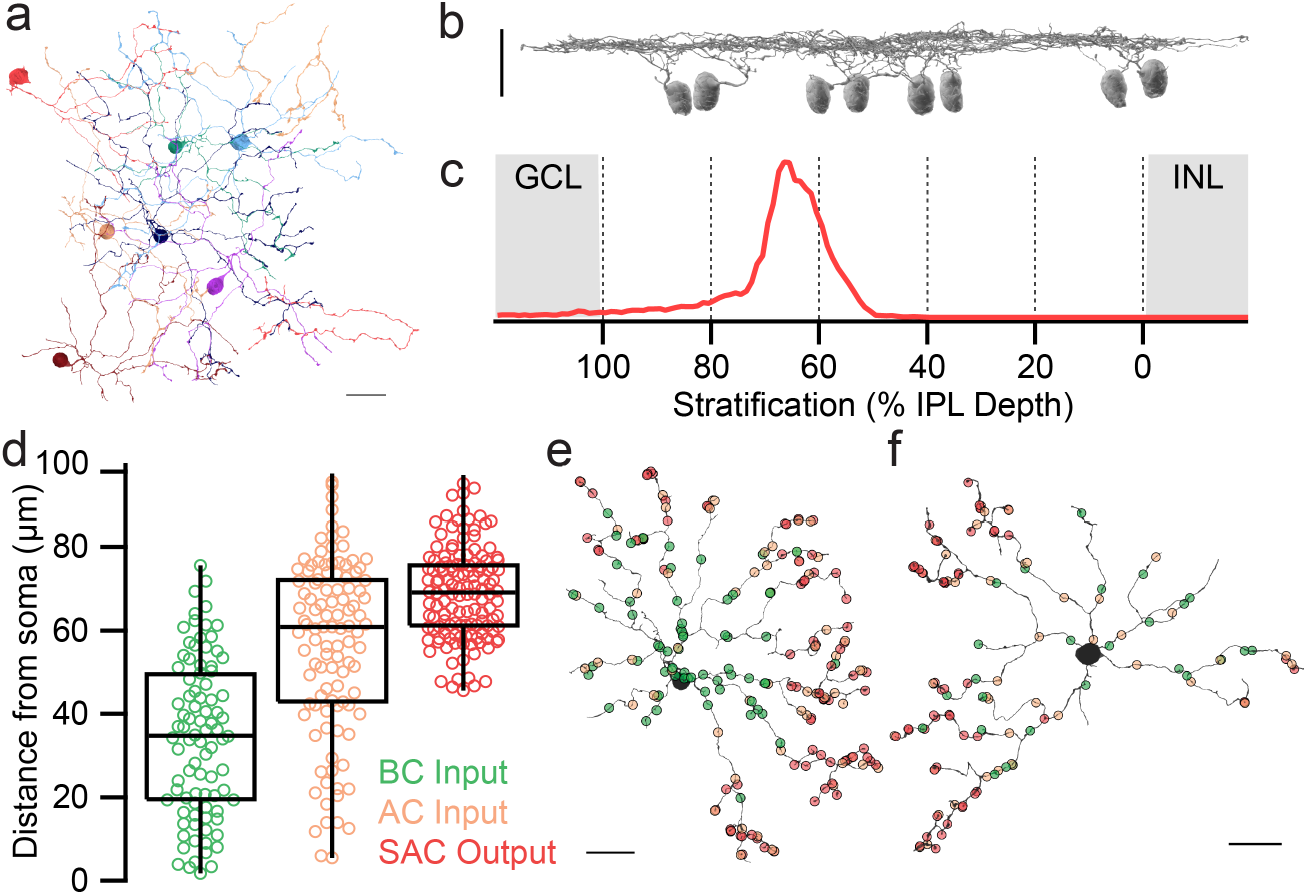
ON SACs of the primate retina. **a,** 3D reconstructions of representative ON SACs and isolated ON SAC dendrites. **b,** Side view of 8 ON SAC reconstructions. **c,** ON SAC stratification depth in the inner plexiform layer (IPL), 67.6 ± 11.0% (mean ± s.d., n = 8 SACs). **d,** Distance from the soma for each synapse type from 2 ON SACs. **e-f,** Locations of ribbon input, conventional input and output synapses along two ON SAC’s dendritic fields. All scale bars are 20 μm.

Direction selectivity in dsRGCs depends critically on SACs^11^. To determine whether primate ON SACs are well-suited to play a similar role, we investigated the structural basis for two features essential to this role: centrifugal motion tuning and asymmetric inhibition of post-synaptic neurons^6^. SACs are ideal for connectomic analysis as decades of careful study in other species has detailed the incredibly precise relationships between their anatomy, physiology and function^6^. We began with the SACs’ selectivity for outward motion, which is supported by cell-intrinsic mechanisms (morphology and radial synapse distribution^22,23^) and amplified by circuit-level mechanisms (temporally-diverse excitatory bipolar cell input^24,25^ and SAC-SAC lateral inhibition)^26,27^. While the relative contributions of these and other mechanisms remains an active area of investigation^2,6^, together they provide a solid blueprint to begin our connectomic investigation. If primate SACs confer direction selectivity on downstream RGCs, we expect to find evidence for these mechanisms.

We first asked whether the proximal-distal synaptic gradient supporting direction selectivity in mammalian SAC dendrites is also present in primates. As expected, the SACs’ output synapses were confined to varicosities on the distal dendrites while bipolar cell input was located closer to the soma, often at the small spines extending from the thin proximal dendrites (**Fig. 1d-f**). This distribution of excitatory input and synaptic output, combined with the SAC’s characteristic morphology, is critical for centrifugal motion tuning^22,23^.

SACs receive the majority of their inhibitory input from neighboring SACs^22^ and the resulting lateral inhibition reduces sensitivity to inward motion^26,27^. We were able to reliably classified the pre-synaptic amacrine cells as SACs or non-SACs for 54 of 63 inhibitory synaptic inputs to the SAC in **Fig. 1e** and confirm that 60.38% came from other ON SACs. Interestingly, the exact location of inhibitory input along each SAC dendrite varies between species and is hypothesized to scale with eye size to preserve angular velocity tuning^22^. SACs in species with larger eyes have greater inter-soma distances and receive inhibition from other SACs more distally. The macaque eye size of 200 μm/degree of visual angle^28^ is consistent with the low SAC coverage factor^5^ and our observed bias of inhibitory synaptic input to the distal dendrites (**Fig. 1a, 1e-f**). These results demonstrate SAC-SAC lateral inhibition is present in primates and positioned to compute a behaviorally-relevant measure of motion direction.

Lastly, we investigated the “space-time wiring” hypothesis^24^, which proposes that sustained bipolar cell input is located closer to the soma than transient bipolar cell input. The resulting temporally-diverse excitation sums for centrifugal, but not centripetal, motion^24,25^; although this mechanism remains controversial^6,22,23^. We reconstructed the axon terminals of presynaptic bipolar cells and classified each as either ON midget, DB4 diffuse, or DB5 diffuse (**Fig. 2a, 2d-f**). The “space-time wiring” hypothesis predicts that midget bipolar cells will be closest to the soma as their responses are more sustained than diffuse bipolar cells^29^. To test this prediction, we calculated the radial distance from the SAC’s soma to each bipolar cell synaptic input. Indeed, ON midget bipolar cells were significantly closer to the soma than the diffuse bipolar cells (**Fig. 2b-c**). However, the underlying distributions show substantial overlap between the two groups. Thus, our results mirror those in other species – while we find evidence for space-time wiring, the overall efficacy could be reduced by the lack of spatial segregation between bipolar cell types^22^.

**Figure 2.**
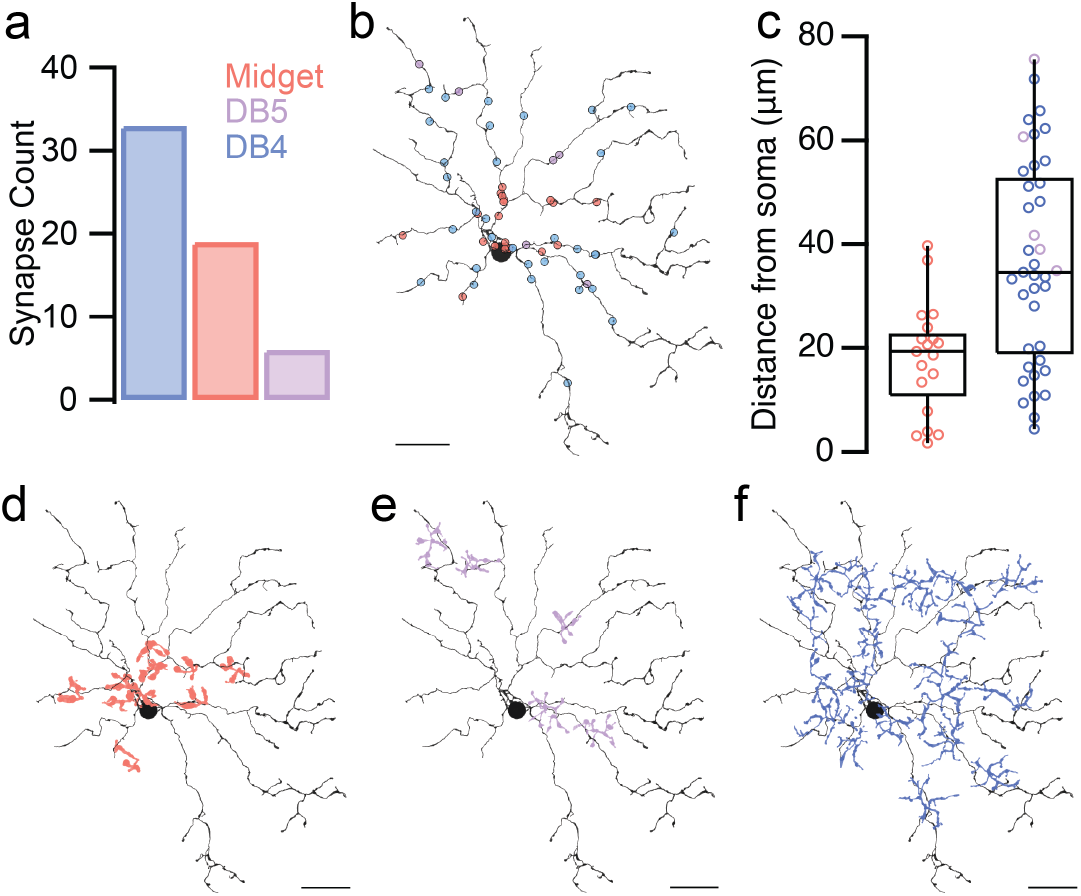
Bipolar input to ON SACs. **a,** Frequency of bipolar cell types presynaptic to ON SACs. **b,** Locations of ribbon synaptic input to an ON SAC, colored by bipolar cell type. **c,** Distance from the soma for ON midget and diffuse (DB4 and DB5) bipolar cells (17.94 ± 2.47 μm vs. 36.72 ± 3.17 μm, mean ± s.d., p = 0.0011). **d-f,** 3D reconstructions of presynaptic axon terminals of ON midget, DB4 diffuse and DB5 diffuse bipolar cells, respectively. All scale bars are 20 μm.

Taken together, our investigation of the structure and synaptic input to ON SACs revealed that multiple mechanisms contributing to centrifugal motion tuning in mammalian SACs are conserved in primates. However, the SAC’s ability to confer direction selectivity depends not only on their responses to outward motion, but also their selective wiring with specific dsRGC subtypes. To address this, we next asked which RGCs were post-synaptic to the ON SACs and whether they received asymmetric SAC inhibition.

## SAC Synaptic Output to RGCs

SAC output synapses have a highly distinctive ultrastructure with large synaptic contacts where the SAC’s processes completely engulf the postsynaptic dendrite^3,21,30,31^. We frequently observed these stereotyped ”wrap-around” synapses at the varicosities of distal SAC dendrites (**Fig. 3b**). We mapped each output synapse, then reconstructed and classified the postsynaptic neurons (**Fig. 3a**). As expected, the SACs’ output targeted other SACs and non-SAC amacrine cells, but rarely bipolar cells. Crucially, RGCs received the majority of the SACs’ output.

**Figure 3.**
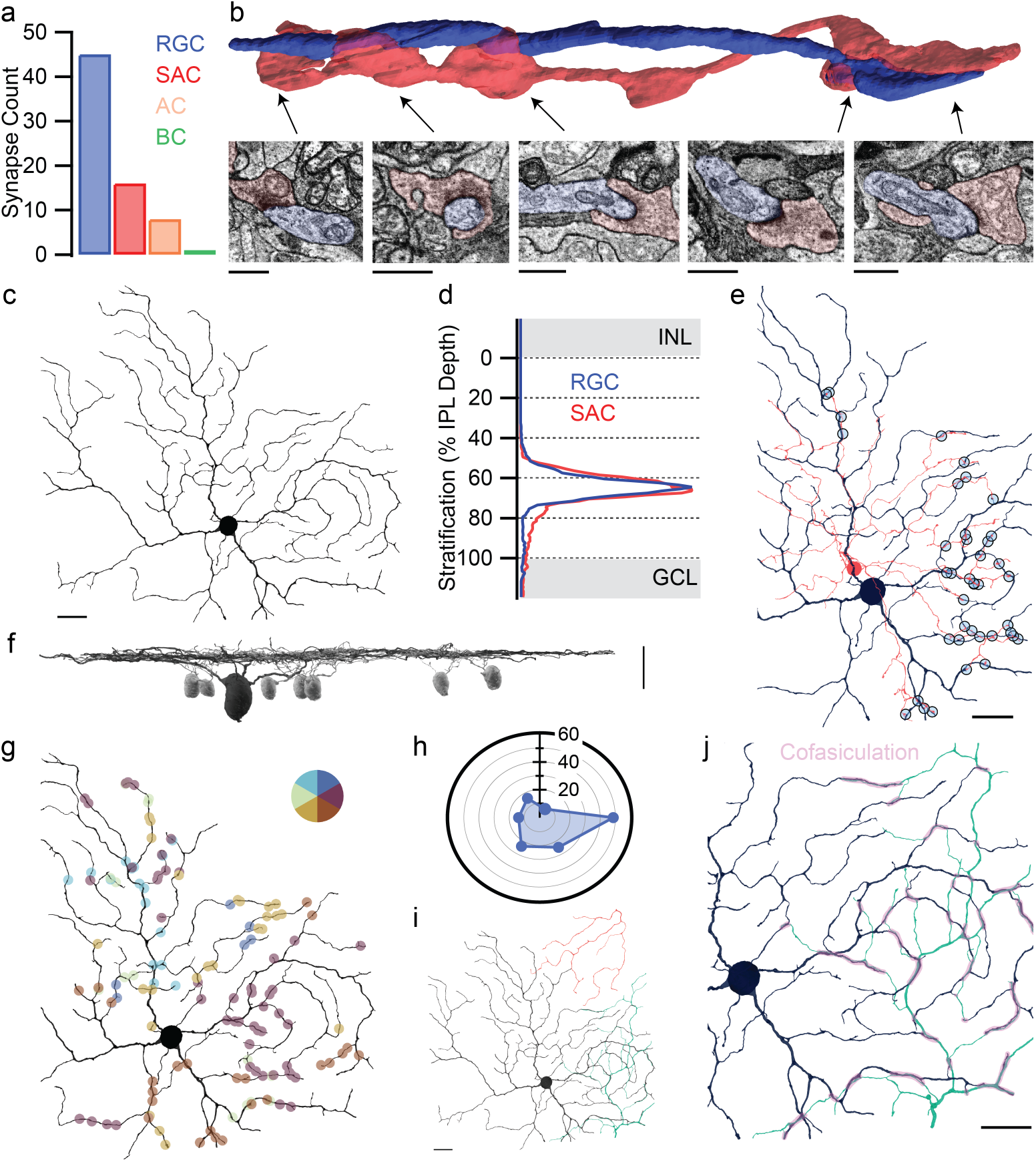
ON SAC synaptic output to RGCs. **a,** Frequency of SAC output to RGCs, SACs, non-SAC amacrine cells and bipolar cells. **b,** 3D reconstruction of representative “wrap-around” synapses between an ON SAC and RGC dendrites. EM micrographs show five distinct synapses between the two cells. **c,** 3D reconstruction of the RGC in **3a-b**. **d,** Co-stratification profile of the rmRGC with the ON SACs from **1b**. **e,** Light blue markers show locations of synapses between the ON SAC and RGC. Note the co-fasciculation between the two cells. **f,** Side view of the RGC in **3c** with ON SACs (light gray). **g,** Locations of SAC synaptic input to the rmRGC, colored by dendritic angle. **h,** Polar histogram of dendritic angles for the synapses depicted in **3g**. **i,** Population of three rmRGCs. **j,** Co-fasciculation of overlapping rmRGCs. Scale bars in **3b** are 1 μm, the rest are 20 μm.

Amazingly, we found the ON SAC’s synaptic output targeted a single RGC type with striking selectivity (**Fig. 3f**). The RGC’s morphology matches the recursive monostratified RGC (rmRGC), a rarely-encountered neuron described only in the largest anatomical surveys of primate RGCs^18,32^. Although the rmRGC’s physiology and connectivity are unknown, a role in direction selectivity has been proposed before on the basis of their strong resemblance to the highly stereotyped morphology of ON-sustained dsRGCs in other vertebrates^33–35^. The recursive, looping branching pattern that ON dsRGCs share with rmRGCs is attributed to their co-fasciculation with the SAC plexus^36,37^. We observed similar co-stratification and co-fasciculation between the rmRGC and ON SACs (**Fig. 3d-e**).

Because each SAC dendrite is independently tuned to outward motion, the directionality of SAC inhibition can be predicted from the angle of the vector between the soma and each output synapse^3,31,38,39^. To estimate the rmRGC’s direction selectivity, we calculated this for **135** of the rmRGC’s conventional synaptic inputs where a presynaptic SAC could be sufficiently reconstructed (**Fig. 3g**). Because the SAC’s output is inhibitory, the rmRGC is predicted to prefer motion in the opposite direction of the strong bias shown in **Fig. 3h**. The distribution of dendritic angles was highly non-uniform (p < 0.0001; Rayleigh test), indicating the asymmetric SAC inhibition necessary for direction selectivity is present.

To confirm our findings, we next searched for additional rmRGCs. The dendritic fields of ON dsRGCs tuned to the same direction tile while those tuned to different directions overlap^39,40^, predicting additional rmRGCs should be present within the first rmRGC’s dendritic field. Indeed, we found additional rmRGCs, both in the first volume taken from the inferior retina and in a separate volume of central nasal retina, each exhibiting the same characteristic morphology, stratification and stereotyped SAC input (**Fig. 3i, S1**). The dendrites of overlapping rmRGCs were often directly adjacent, consistent with mammalian ON dsRGCs’ cofasciculation with each other and the ON SAC plexus^39,40^ (**Fig. 3i-j**). Moreover, puncta adherens were frequently observed between adjacent rmRGC dendrites, a unique feature previously reported in rabbit ON dsRGCs^30^ (**Fig. S2**).

While the other rmRGCs’ proximity to the volume’s edge prevented an unbiased dendritic angle analysis, we did observe a striking trend: the SAC dendrites providing input to the first rmRGC rarely, if ever, synapsed on the second rmRGC in **Fig. 3j**, despite frequently being in close proximity. For example, the SAC in **Fig. 3e** synapsed on the first rmRGC 40 times, but only three times on the second rmRGC (**Fig. S3**). These results are consistent with the hypothesis that the overlapping rmRGCs prefer movement in different directions and supports asymmetric inhibition from SACs as an underlying mechanism for their distinct directional tuning. Taken together, the rmRGC’s structure and circuitry is consistent with ON dsRGCs in other species.

## Retinal Input to the AOS

Like SACs, ON dsRGCs play a fundamental and highly conserved role in vision^41^. From mice and rabbits to turtles and birds, ON dsRGCs share not only the characteristic morphology shown in **Fig. 3c**, but also projections to the AOS, which coordinates with the vestibular system to stabilize gaze with the optokinetic reflex^34,35,42,43^. As in other species, neurons in the primate AOS exhibit similar response properties to ON dsRGCs^42,44^ and receive direct retinal input^45^, yet the specific RGC types are unknown. Previous retrograde tracer injections to the nucleus of the optic tract and dorsal terminal nucleus (NOT-DTN) complex of the AOS revealed input from two RGC types, one resembling the rmRGC^46^; however, incompletely-filled dendritic fields prevented unambiguous classification. We repeated this experiment with the goal of targeting individual RGCs for detailed morphological characterization and comparison to our reconstructions.

We injected the retrograde tracer rhodamine dextran into the NOT-DTN^47^, which was located by identifying neurons with characteristic response properties, including direction selectivity during horizontal smooth pursuit (**Fig. 4a, S4a**). Further confirmation was obtained with post-mortem histology (**Fig. S4b**). Retrogradely-labeled RGCs were identified in an *ex vivo* flatmount preparation by clumps of rhodamine fluorescence within their soma (**Fig. 4b**) and filled with fluorescent dye to reveal their dendritic field structure (**Fig. 4c-d, S5**). All were rare wide-field RGCs and a subset exhibited the characteristic curving dendrites that are hallmarks of both rmRGCs and ON dsRGCs, confirming the NOT-DTN of the AOS receives direct rmRGC input.

**Figure 4.**
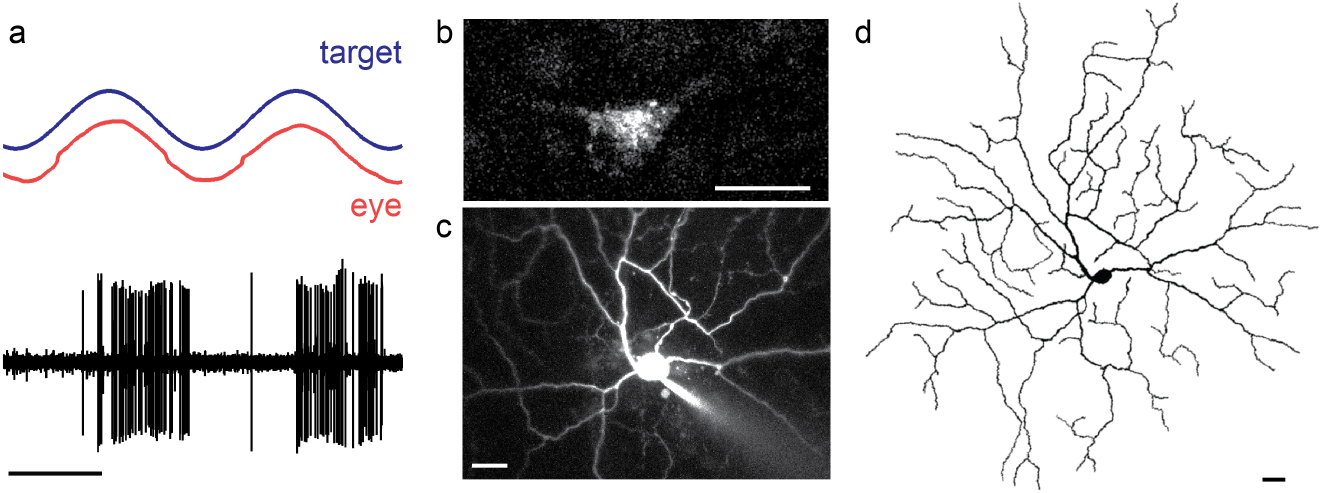
Retrograde labeling of RGCs projecting to the NOT-DTN. **a,** Direction-selective response properties recorded from the NOT-DTN to horizontal smooth pursuit. Scale bar is 1 second. **b**, RGC soma identified by fluorescent rhodamine dextran granules. **c**, Cell fill of rhodamine-labeled RGC, electrode to the bottom left. **d**, Tracing of the dendritic field in **4c**, omitting the axon. Scale bars in **b-d** are 20 μm.

## Discussion

Here, we identified a likely homolog to the ON dsRGC, establishing an anatomically-conserved circuit from SACs to the AOS for the gaze-stabilizing, compensatory eye movements essential for visually-guided navigation^42^. Past research on the primate AOS has largely focused on the role of cortical feedback; however, recent studies on human congenital nystagmus suggest a direct retinal contribution involving GABAergic signaling by SACs^13,14,48^. Until now, the RGCs linking SACs to the AOS were unknown.

We estimate that rmRGCs make up ~1% of all RGCs in the central retina. This rarity likely reflects their large dendritic fields (**Fig. S6**) rather than their importance in vision. The low density of rmRGCs undoubtedly creates a challenging sampling bias that could explain their absence in surveys recording from single RGCs with microelectrodes or relatively small patches of peripheral retina with multielectrode arrays. Although follow-up physiological studies guided by our results will be important, waiting for these experiments to become feasible delays essential insights into primate vision. Here we demonstrate that connectomics – a technique best known for large-scale dense reconstructions^24,39,49^ – can also be utilized for focused, hypothesis-driven questions about otherwise intractable neural circuits^19^ with implications for human health and disease^13,14,48^.

Interestingly, we did not observe SAC input to a ON-OFF dsRGC homolog, although we cannot rule out the possibility that they are present in the peripheral retina where OFF SAC density increases. However, the central retina mediates most conscious vision, indicating any ON-OFF dsRGC homolog confined to the periphery would be best suited for non-image-forming vision and independent of the cortical direction selectivity underlying motion perception^50^.

This work underscores the benefits of bridging primate and non-primate research^16^. The SAC and rmRGC join a growing list of primate retinal neurons, like the intrinsically photosensitive RGCs, that are understood only through prior research in non-primate species. The strong correspondence between rmRGCs and ON dsRGCs raises the intriguing possibility that other rare primate RGCs with unknown functions may have well-studied non-primate counterparts. More generally, this challenges a widely-held view that the computational goals of the primate retina are unique from other species and maximize information transmission to the cortex rather than extraction of meaningful visual features, such as motion direction.

## Acknowledgements

We are grateful to James Kuchenbecker and James Anderson for IT support, Andrea Bordt and Rebecca Girresch for annotation assistance and Chris Chen and Michelle Giarmarco for assistance with tissue preparation. We also thank Shellee Cunnington, Ed Parker and Jessica Rowlan for excellent technical assistance. Tissue was provided by the Tissue Distribution Program at the Washington National Primate Research Center with the help of Chris English. This work was supported by NIH grants R01-EY027859 (J.N.), R01-EY028902 (R.S.), F32-EY032318 (S.P), T32-EY007125 (S.P), T32-EY007031 (S.P), P30-EY014800, P51-OD010425 and Research to Prevent Blindness. Implementation of the Viking software environment for image capture, registration, database, and annotation was supported through NIH grants to Bryan William Jones (R01-EY015128, R01-EY028927).

## Author Contributions

S.P. conceived of the project, wrote the paper and performed the serial EM experiments. J.N., S.P., F.R. and R.S. designed experiments. J.N. edited the paper. J.N and M.N acquired the serial EM volume. B.B. and R.S. performed the retrograde tracer injections. F.R. and S.P. performed the cell fills. B.B., M.M. and S.P performed the immunohistochemistry and confocal microscopy.

## SUPPLEMENTARY FIGURES

**Figure S1.**
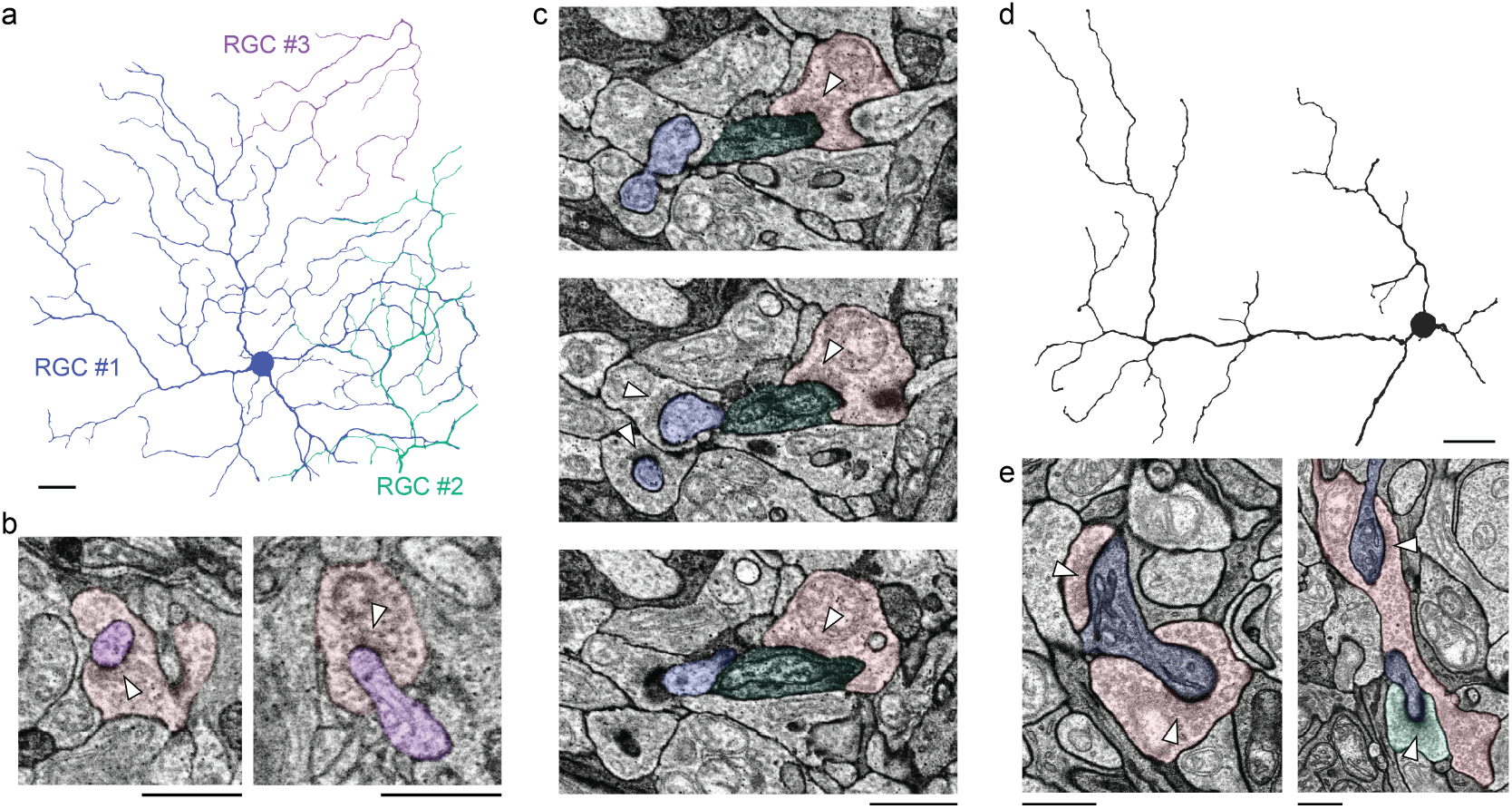
Additional rmRGCs with characteristic “wrap-around” SAC synaptic input. **a,** Dendritic fields of three rmRGCs in the first serial EM volume. **b,** EM micrographs of SAC (pink) input to rmRGC #3 (purple). **c,** EM micrograph series of SAC input to rmRGC #2 (green). Note input from other SACs to nearby rmRGC #1 (blue). **d,** Partial reconstruction of an rmRGC from a second serial EM volume of nasal retina. **e,** EM micrographs of the characteristic “wrap-around” synapses from SACs (pink, green) onto the rmRGC (blue). Scale bars in **a** and **d** are 20 μm, the rest are 1 μm.

**Figure S2.**
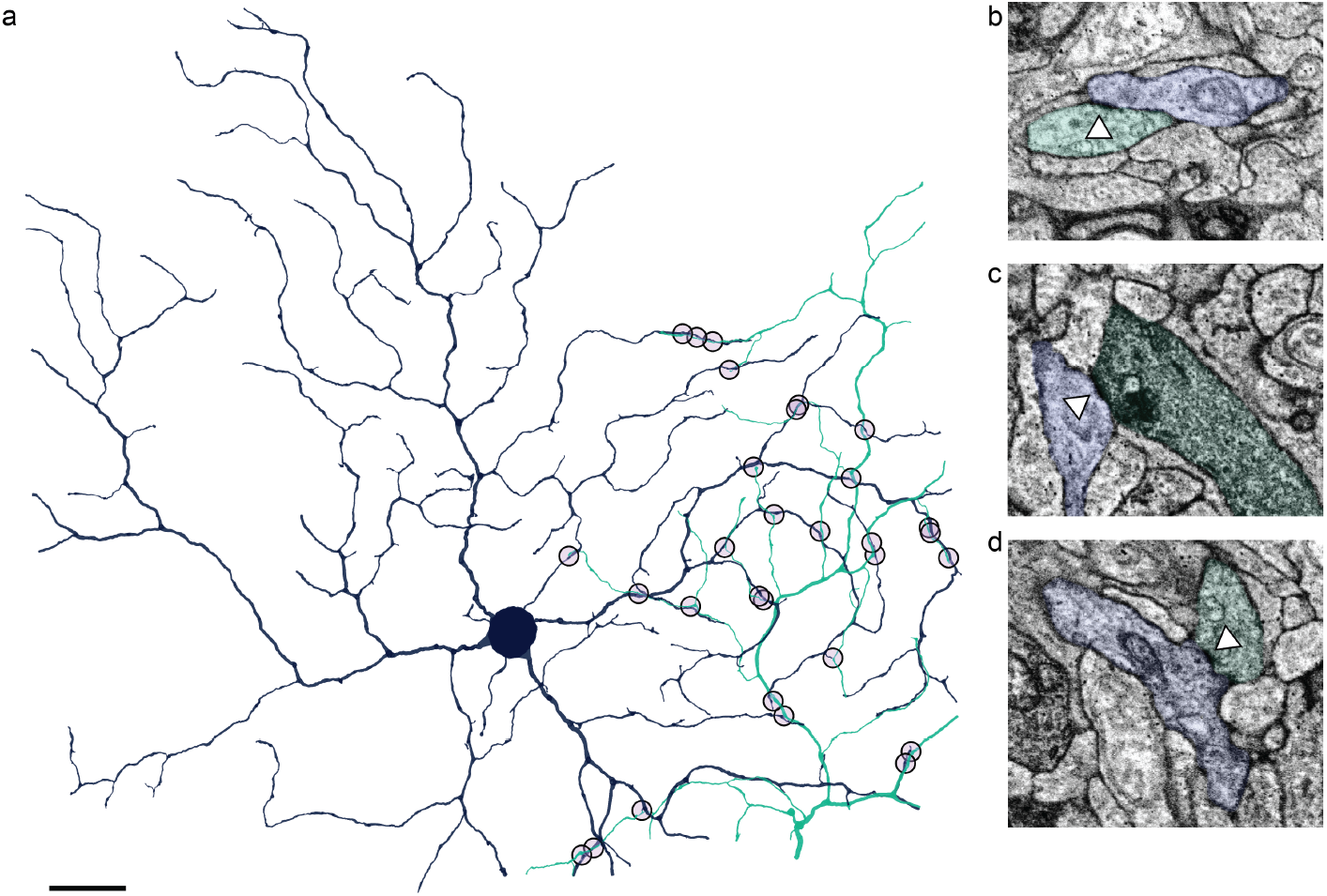
Additional identifying features of recursive monostratified RGCs. **a,** Markers indicate the locations of all puncta adherens between the two primary rmRGCs in **Fig. 4j**. Scale bar is 20 μm. **b-d,** Representative EM micrographs of puncta adherens, symmetric membrane densities without synaptic specialization^30^.

**Figure S3.**
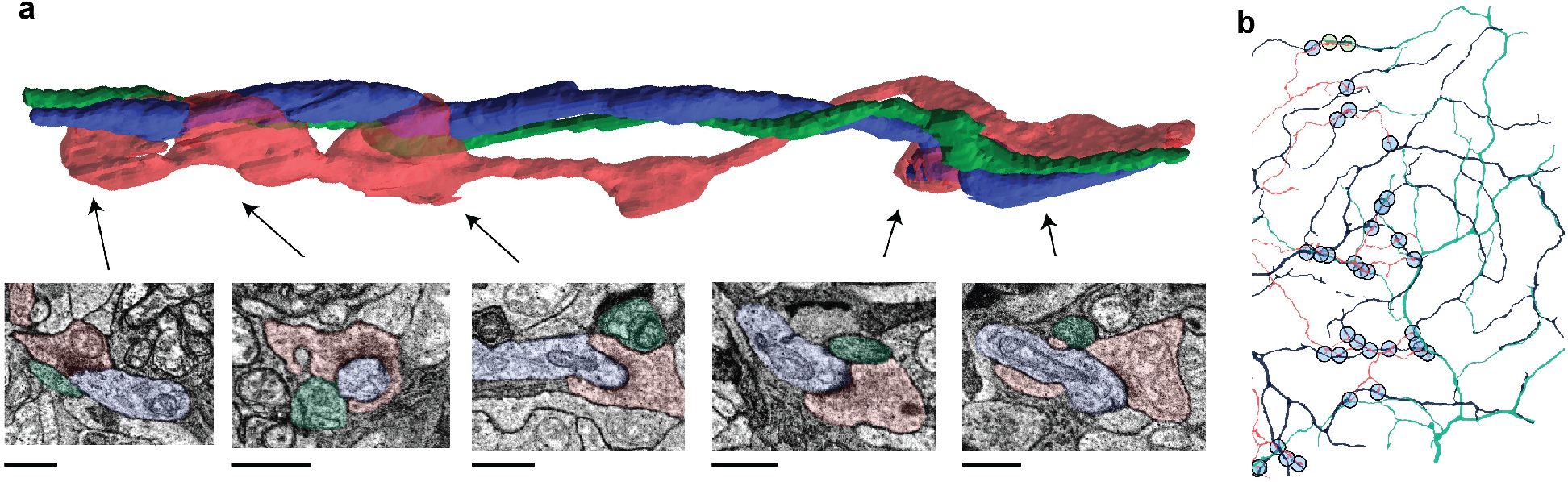
Lack of shared SAC input between overlapping rmRGCs. **a,** As in **Fig. 3b**, but with the 2^nd^ rmRGC from **Fig. 3j** added in green. Note the close proximity to the first rmRGC and SAC, but lack of input from the SAC. Scale bars are 1 μm. **b,** As in **Fig. 3e**, but with synapses onto the 2^nd^ rmRGC marked in green.

**Figure S4.**
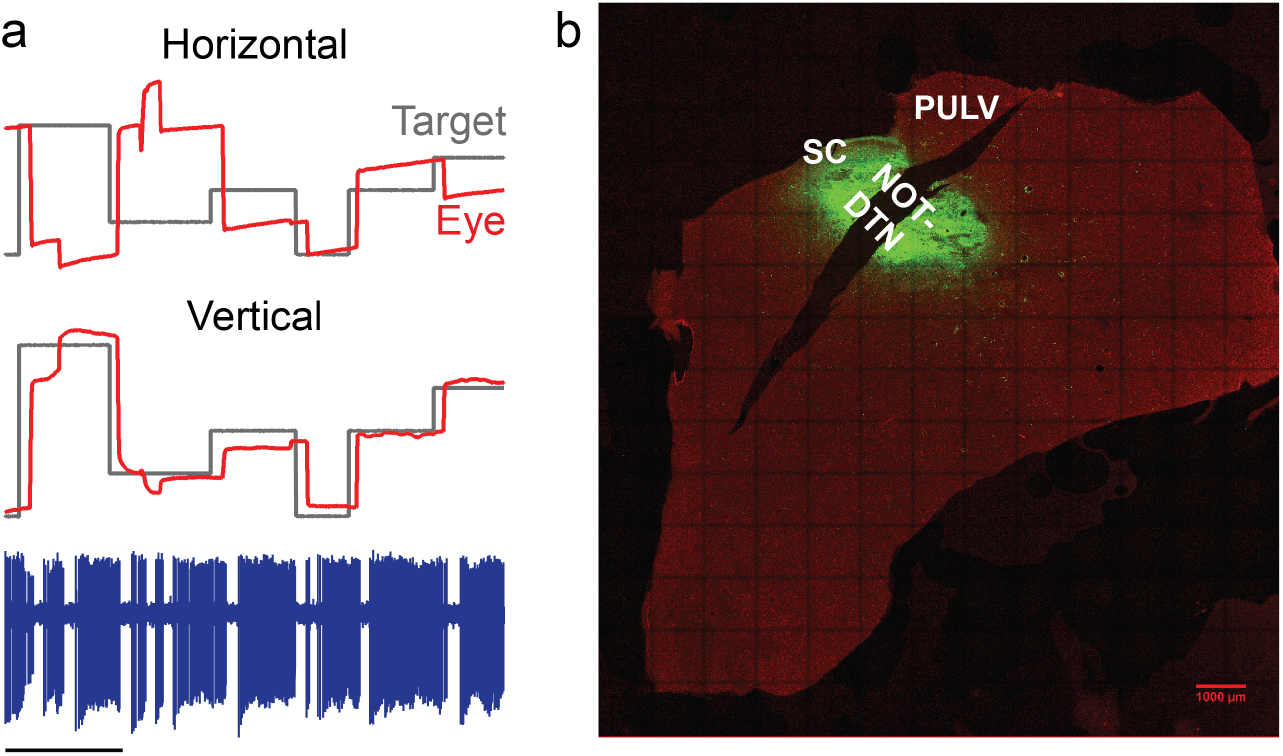
Additional verification of NOT-DTN injection site. **a,** FOPN recorded while confirming the boundaries of the NOT-DTN, adjacent to the area where directionally-selective neurons were encountered. Scale bar is 1 second. **b,** Injection site. Rhodamine fluorescence (green) marks the injection site relative to the left superior colliculus (SC) and pulvinar (PULV), DAPI (red) labels all nuclei. Scale bar is 1 mm.

**Figure S5.**
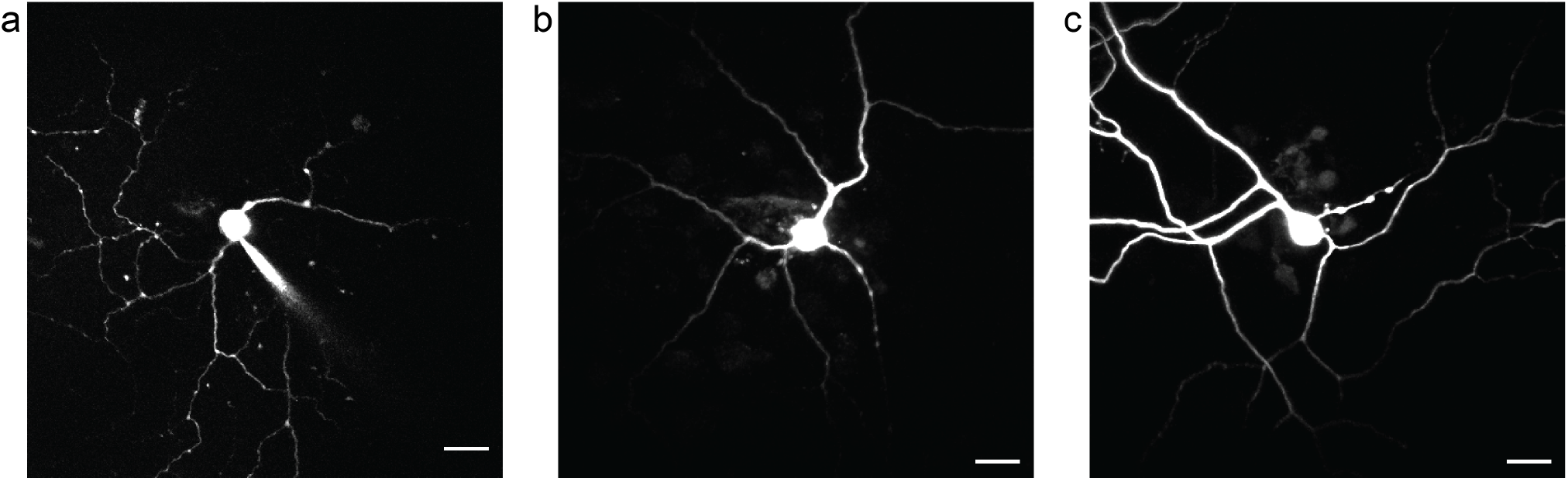
Additional retrogradely-labeled RGCs. **a,** Another recursive monostratified RGC. **b-c,** Two unidentified wide-field RGCs. All scale bars are 20 μm.

**Figure S6.**
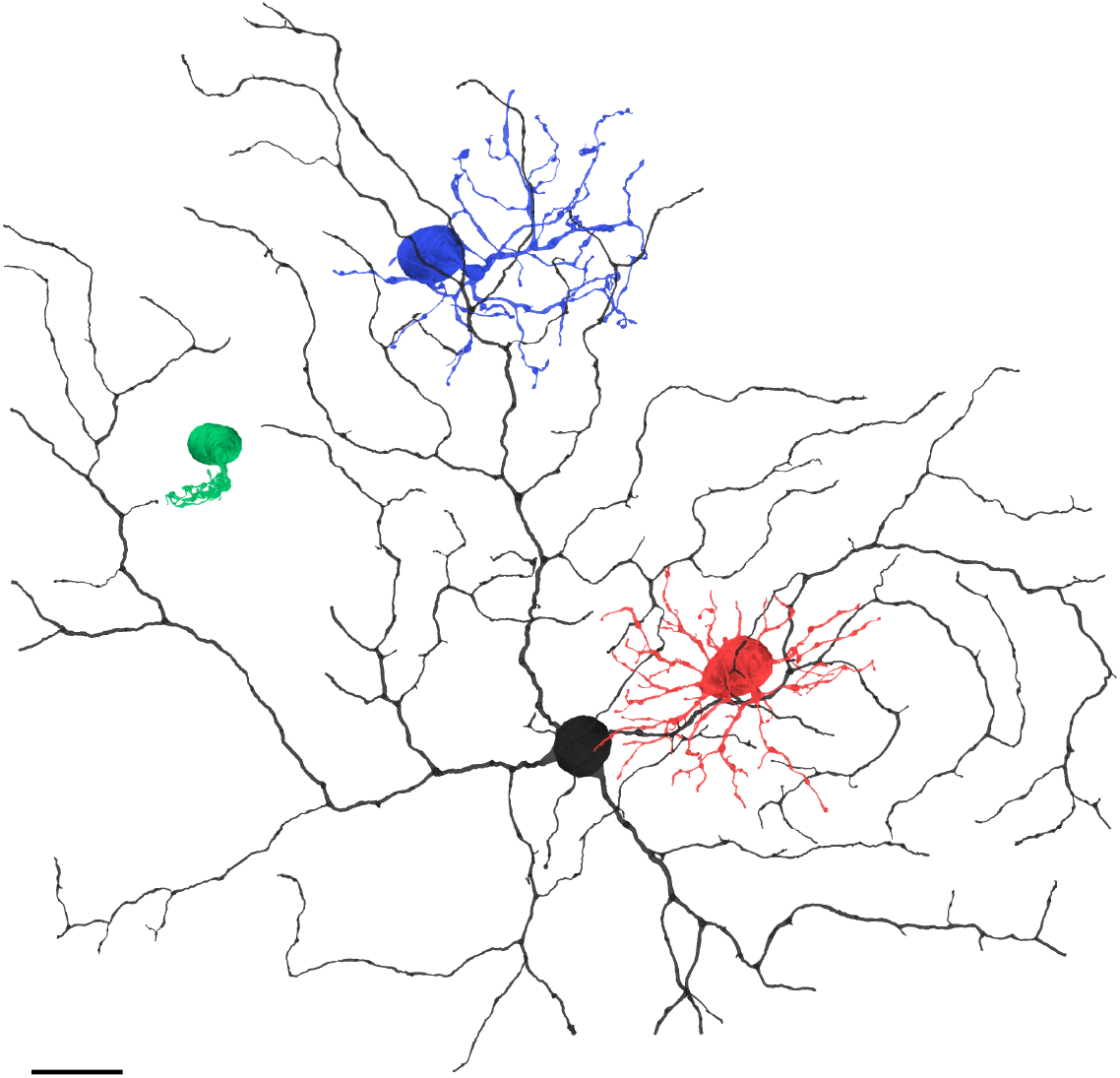
Comparison of rmRGC dendritic field with other common primate RGCs. **a,** An rmRGC dendritic field (black) compared to the three most common and well-studied primate RGCs: parasol (red), small bistratified (blue) and midget (green). Scale bar is 20 μm.

## METHODS

### Serial Electron Microscopy

Retinal tissue for serial electron microscopy was obtained from a terminally anesthetized male macaque (*Macaca nemestrina*) monkey though the Tissue Distribution Program at the Washington National Primate Center. All procedures were approved by the Institutional Animal Care and Use Committee at the University of Washington. Blocks of inferior and nasal parafoveal retinal tissue at ~1 mm eccentricity from the foveal center were processed as previously described^51^. The inferior retinal tissue was used for all results other than **Fig. S1d-e**. A cross-section of the inferior retinal tissue taken with a transmission electron microscope can be found in our previously published work^52^.

The tissue was imaged using a Zeiss Sigma VP field emission scanning electron microscope equipped with a 3View system and sectioned in the horizontal plane. Tissue preparation and image collection were optimized in signal-to-noise ratio for visualizing small, low contrast features such as synaptic ribbons that have previously been a challenge for serial block-face scanning electron microscopy. The volume of inferior retina was imaged at a resolution of 7.5 nm/pixel and contained 1893 sections at 90 nm section widths, spanning from the ganglion cell layer (GCL) through the cone pedicles. The volume of nasal retina was imaged at a resolution of 5 nm/pixel and contained 2355 sections at 50 nm section widths, spanning from the GCL to the inner nuclear layer (INL). While the size of the volume limited the number and dendritic field extent of rmRGCs and SACs that could be analyzed in detail, our reconstructions are comparable to other recent serial EM studies^3,22,38^.

### Annotation

Initial image registration was performed with Nornir (RRID: SCR_003584, http://nornir.github.io) and supplemented where necessary with custom MATLAB (Mathworks) code or the SIFT feature registration plugin for ImageJ. The detailed reconstructions in **Fig. 3b** and **S3a** were annotated by contouring the neuronal processes using TrakEM2 (RRID:SCR_008954, http://www.ini.uzh.ch/~acardona/trakem2.html)^55^. All other annotation was performed with Viking (SCR_005986, https://connectomes.utah.edu^53^. Neuronal processes were reconstructed through the sections by placing a circular disc at the structure’s center of mass and linking the disc to annotations from the same structure on neighboring sections. Synapses were annotated with lines connected by 2-3 control points and linked to a parent neuron. The synapse annotations for the presynaptic and postsynaptic neurons were also linked to each other so that the annotated neurons and the specific links between them could be represented and queried as a network.

We used established criteria for identifying synapses^54^ and confirmed that the synaptic structures spanned more than one section. Conventional synapses were identified by a cluster of vesicles within the presynaptic neuron adjacent to a membrane density on the post-synaptic neuron. Ribbon synapses were identified by the presence of a ribbon structure adjacent to a membrane density on the post-synaptic neuron. The puncta adherens between rmRGC dendrites were identified as symmetrical membrane densities without any evidence of specialization for synaptic transmission^30^. These contacts were unlikely to be gap junctions because SBFSEM in general and our volumes specifically lack the resolution for gap junctions and none were observed along cell types known to make large gap junctions, such as AII amacrine cells.

### Cell Type Identification and Classification

ON SACs were identified by their highly characteristic morphology, ultrastructure and stratification^5,21,30^. Their somas were displaced to the GCL with a single primary dendrite that split into several secondary dendrites, expanding radially to stratifying in sublamina 4 (**Fig. 1**). The proximal dendrites were extremely thin while distal dendrites were covered in varicosities. To classify pre- and post-synaptic amacrine cells, each neuron was annotated until a confident classification of SAC vs. non-SAC could be made. In addition to the morphological features described above, isolated SAC dendrites were also classified by their synapses and connectivity. For distal dendrites, this included “wrap-around” synapses at varicosities, reciprocal synapses with other SACs and an absence of synaptic output to bipolar cells. For proximal dendrites, criteria included thin dendritic diameters, sparse radial branching, an absence of synaptic output sites and bipolar cell input, often at the end of short dendritic spine-like branches. Dendritic diameter and branching frequency in particular distinguished SACs from other S4 amacrine cells, such as the semilunar and wiry amacrine cells^56^. These morphological features reported previously and confirmed in our full SAC reconstructions served as a guide for confirming isolated branches as SACs^3,22,30,31^.

The rmRGCs were primarily identified by their characteristic curving dendrites. This dendritic field structure is similar to that of the recursive bistratified RGC; however, the recursive bistratified RGC dendrites frequently change strata and, as a result, often overlap when viewed in a flatmount. Overlapping dendrites were rarely observed with the rmRGCs and largely limited to places where distal dendrites travelled under primary dendrites. As reported for ON dsRGCs in other species^39^, we occasionally observed isolated dendrites extending towards the OFF sublamina; however, these branches ended quickly and were too sparse to form a clear second dendritic tier as reported for recursive bistratified RGCs.

We classified the ON bipolar cells presynaptic to SACs and rmRGCs into four types – ON midget, giant, DB4 and DB5 – following previous serial EM classifications^57^. ON midget bipolar cells stratified closest to the GCL and exhibited a small, but densely-packed axon terminal. In several cases, further confirmation was obtained by partially reconstructing the post-synaptic ON midget RGC. Giant bipolar cell axon terminals had sparse branches covering a large area. The distinction between DB4 and DB5 bipolar cells was more subtle as the two costratify and have similar dendritic fields. The branches of DB5 bipolar cell axon terminals were generally smaller, denser and exhibited more varicosities than those of DB4 bipolar cells. In some cases, additional verification was obtained by reconstructing the ON-OFF lateral amacrine cells previously reported to form reciprocal synapses with DB4, but not DB5, bipolar cells^57^.

### Analysis

All analyses on the serial EM data were performed with open source connectomics toolbox, SBFSEM-tools (RRID: SCR_005986, https://github.com/neitzlab/sbfsem-tools), in MATLAB (Mathworks). The Rayleigh test from the CircStat toolbox for MATLAB was used to determine whether the distribution of dendritic angles was uniform or not^60^. In all other cases, the Wilcoxon rank sum test was used to determine statistical significance. The box plots in **Fig. 1d** and **2c** show the maximum, 75^th^ percentile, median, 25^th^ percentile and minimum values.

Stratification depth within in the inner plexiform layer (IPL) was calculated as previously described^56^. Briefly, markers were placed throughout the volume at the borders between the INL-IPL and IPL-GCL. The final INL-IPL and GCL-IPL boundaries were determined by surfaces fit to the X, Y and Z coordinates of the markers denoting each boundary type using bicubic interpolation. Given an annotation’s X and Y coordinates, the surfaces returned the Z-coordinates of both IPL boundaries at that X, Y location. The annotation’s Z coordinate relative to the Z coordinates of each boundary could then be calculated to determine percent IPL depth. In this way, IPL depth was calculated for each annotation individually to account for local variations in IPL thickness and the volume’s slope due to its proximity to the edge of the fovea. The accuracy of this approach is supported by the strong correspondence between the stratification of ON SACs within our volume and values previously reported in the literature^58,59^. For **Fig. 1b** and **3d**, the Z position of each annotation was corrected by normalizing for local IPL variations and translating by the local offset calculated from the INL-IPL and GCL-IPL boundary surfaces.

The radial distance of synapses from each SAC’s soma in **Fig. 1d** and **2c** was calculated as the 2D Euclidean distance. Because each SAC was monostratified, the 2D distance was used rather than the 3D distance to avoid introducing artifacts associated with the slope of the volume discussed above. The soma location was automatically chosen as the center of the largest annotation in each SAC. For synapses annotated across multiple sections, the midpoint annotation was used.

### Visualization

The 3D rendering was performed with SBFSEM-tools, as previously described20. Briefly, the 3D models are triangle meshes built by rendering segments of connected annotations as rotated cylinders centered at each annotations’ XYZ coordinates and scaled by their radii. The exception was the detailed reconstructions in **Fig. 3b** and **S3a**, which were rendered with TrakEM2 instead. Where applicable, RGC axons were omitted to emphasize the dendritic field structure. Figures were prepared in either MATLAB, ImageJ or Igor Pro 8 (RRID:SCR_000325, Wavemetrics) and final layouts were arranged in Adobe Illustrator.

### Retrograde tracer injections

This experiment was performed on a 15.1 kg adult rhesus monkey (*Macaca mulatta*). The NOT-DTN was approached through a chamber targeting the superior colliculus. The chamber was tilted to the left by 20° and aimed at a point in the midline, 9 mm dorsal and 2 mm anterior to stereotaxic zero. The NOT-DTN was identified physiologically by recording from neurons tuned to horizontal pursuit in specific directions (**Fig. 4a**)^44^. Additionally, the following omnidirectional pause neurons (FOPNs) previously reported to be located dorsal to the directionally-selective neurons of the NOT-DTN were identified while approaching the injection site from the foveal superior colliculus (**Fig. S4a**)^61^. The locations of the directionally-selective neurons and FOPNs were used to map the boundaries of the NOT-DTN prior to the injection.

5% biotinylated dextran-conjugated tetramethyl rhodamine 3000 MW (micro ruby, #D-7162; Molecular Probes, Eugene, OR) in distilled water^47^. Three injections of 250 nL, 480 nL and 560 nL were made distanced ~300 μm apart. The first injection site was where the first FOPN was encountered and the final injection site was where background activity corresponding to smooth pursuit was heard.

To visualize the rhodamine fluorescence marking the NOT-DTN injection site, the brain was fixed by perfusing the animal with 4% paraformaldehyde in 0.1 M phosphate buffer. After the brain was removed from the skull, it was brought up in 30% sucrose in 0.1 M phosphate buffer. The injected rhodamine dextran could be seen in the fixed brain as bright pink and the surrounding thalamus and midbrain were cryosectioned.

### Cell fills

Retinal tissue was prepared as previously described^62^. Briefly, enucleated eyes were hemisected and the vitreous humor was removed mechanically. When necessary, the eye cup was treated for ∼15 minutes with human plasmin (∼50 μg/mL, Sigma or Haematologic Technologies) to aid vitreous removal. Small pieces of the retinal tissue were detached from the retinal pigment epithelium and placed onto the stage of an electrophysiology rig ganglion cell side up. The tissue was superfused with warmed (32-35°C) Ames’ medium (Sigma). All cell bodies were visualized with a 60x objective under infrared illumination. RGCs retrogradely labeled with rhodamine dextran were identified with either one- or two-photon microscopes, then filled with two fluorescent dyes: Alexa-488 to image immediately and Lucifer Yellow for imaging after immunohistochemistry. Afterwards, the retinal tissue was detached from the retinal pigment epithelium if necessary and immersion-fixed in 4% paraformaldehyde in 0.1M phosphate buffer (PB), pH 7.4, for 30 minutes at room temperature, then washed in PB.

### Immunohistochemistry and confocal microscopy

Retinal tissue and brain sections were mounted on glass slides using DAPI Fluoromount-G (SouthernBiotech, 0100-20). Confocal Z-stacks of the rhodamine dextran in retrogradely-labeled RGCs, Lucifer Yellow-filled RGCs and DAPI-labeled nuclei were taken with a Leica TCS SB8 microscope. A 20x oil-immersion objective was used for retinal tissue and a 10x objective for brain slices. For final figure display, ImageJ was used to converted Z-stacks to maximum intensity Z-projections and, in some cases, remove shot noise with the “despeckle” function. The dendritic field in **Fig. 4d** was traced from the original highest resolution Z-stack, using the Simple Neurite Tracer ImageJ plugin^63^.

## Data and Code Availability

The Viking software for visualizing the dataset and the annotations is freely available (https://connectomes.utah.edu). The SBFSEM-tools software used to analyze and render the annotations is open source (https://github.com/neitzlab/sbfsem-tools). All data generated and/or analyzed during the current study are available from the corresponding author on reasonable request.

